# Inflammatory Agonists Modulate the Host Response to Type 2 ECM scaffold Immune Environment and Long-Term Remodeling After Severe Traumatic Injury

**DOI:** 10.1101/2025.11.09.687444

**Authors:** Weizhen Li, Iris Baurceanu, Sanjay Pal, Rohan Chaudhari, Adrienne E. Kimmel, Matthew T. Wolf

**Affiliations:** Cancer Biomaterials Engineering Section, Cancer Innovation Laboratory, Center for Cancer Research, National Cancer Institute, Frederick, MD 21702, USA; Cancer Innovation Laboratory, Center for Cancer Research, National Cancer Institute, Frederick, MD 21702, USA; Department of Cell, Developmental and Cancer Biology & Knight Cancer Institute, Oregon Health and Science University, Portland, OR 97201, USA

## Abstract

The immune system is a vital regulator of tissue repair after trauma and the response to implantable scaffolds for regenerative medicine. Decellularized extracellular matrix (ECM) scaffolds promote tissue integration and remodeling following traumatic injury in part due initiating a pro-reparative Type-2 immune response. However, exogenous soluble inflammatory immune signals can be introduced during scaffold implantation, including microbial products in contaminated surgical fields or during immunotherapy to treat autoimmunity and cancer. It remains largely unknown how such immune mediators modulate the ECM scaffold immune environment and subsequent scaffold remodeling. In the present study, we co-delivered 3 distinct inflammatory immune adjuvants (cyclic di-AMP [CDA], monophosphoryl lipid A [MPLA], and granulocyte colony stimulating factor [GM-CSF]) with small intestinal submucosa (SIS) ECM in a murine volumetric muscle loss injury model, evaluating acute (1 week) and long-term (8-week) immune environments and scaffold remodeling. High parameter spectral cytometry, histologic analysis, and PCR revealed differential potentiation of the ECM scaffold microenvironment. Type 2 immune programs including IL-4, eosinophils, CD4 T cells, and CD206/CD86 macrophage ratios were induced in all ECM groups but attenuated by varying amounts with the CDA and MPLA co-delivery. By 8-weeks, inflammation had largely subsided, and histologically ECM with GM-CSF or MPLA showed the greatest degradation and remodeling into adipose tissue. These findings suggest that early pro-inflammatory compound delivery does not abrogate the ECM immune environment but does attenuate some programs while inducing others. These have long-term effects on scaffold remodeling and should be a consideration for surgical reconstruction in patients receiving immune stimulatory therapies.

## 1. Introduction

Extracellular matrix (ECM) scaffolds prepared from decellularized tissues are immunomodulatory biomaterials for musculoskeletal tissue engineering. When implanted in a site of injury, these acellular scaffolds provide a template to guide host cell infiltration and subsequent remodeling into new tissue as the scaffold is degraded. Successes in clinical applications such abdominal wall reconstruction have prompted investigation into implementing ECM scaffolds in more challenging repairs such as volumetric muscle loss (VML) where the injury exceeds muscle reparative capacity leading to permanent loss of function^1^. Thus, three-dimensional volume filling scaffolds that can bridge these critical sized deficits and permit functional restoration are necessary for these complex cases. The tissue decellularization process is central to ECM scaffold preparation, and when performed appropriately, removes cells, while leaving the ECM largely intact. This ECM is well conserved among mammalian species and provides structural support to infiltrating cells, a reservoir of adhesion ligands, and matrix-bound signaling molecules that directly influence cell behavior during tissue repair^2^. Further, the removal of cell membrane and major histocompatibility complexes eliminates acute tissue rejection to allow allo- and xeno-geneic ECM sources. Although ECM scaffolds do not trigger pro-inflammatory rejection, they are also not immunologically inert but rather generate a Type 2 biased immune environment rapidly after implantation, which directly impacts remodeling outcome.

Immune cells and signals play an indispensable role in both the response to traumatic injury and to implantable materials, and which intersect in musculoskeletal tissue engineering. Skeletal muscle injury classically begins with an inflammatory cascade of blood factors, pro-inflammatory alarmin^3^ release from necrotic myofibers such as HMGB1 and S100 proteins, and phagocyte mobilization (neutrophils and macrophages) to remove damaged muscle^4^. This acute phase is followed by re-polarization to Type 2 immune signals such as M2-like macrophages, eosinophils, and Th2 T cells of adaptive immunity that release cytokines such as interleukin-4 (IL-4) to encourage de novo fibrous tissue formation and myogenesis^5^. ECM scaffolds are unique from other polymeric implants, in that rather than an inflammatory foreign body reaction, ECM biomaterials initiate a short pro-inflammatory phase that quickly resolves to a robust Type 2 immune response. Ablation of specific immune cell types and regulators in preclinical studies have revealed that ECM scaffold remodeling is vitally dependent on M2-like macrophages and IL-4 signaling, which parallels muscle repair. Further, there is evidence that perturbations to initial inflammatory events can have long-term impacts on tissue repair in abrogating the foreign body reaction^6^. Likewise, the temporal sequence of an early Type 1 inflammation proceeding to Type 2 is crucial to repair and can be modulated by immune factors in the scaffold environment.

Exogenous immune signals introduced during implantation surgery are known to interact with the ECM scaffold inflammatory progression. These signals are often derived from microbial products introduced in contaminated surgical fields following open traumatic wounds. Indeed, ECM scaffolds are often implanted in cases that are high-risk of contamination because they are degradable and resistant to biofilm formation. Likewise, lipopolysaccharide (also known as endotoxin) is a toll-like receptor-4 (TLR-4) agonist from gram-negative bacteria and is a known contaminant during material processing. The effect of these signals on repair is mixed. Inactivated bacteria were shown to alternately promote or hinder osteogenesis depending on the model used. Doping endotoxin into ECM scaffolds increased initial inflammation but did not impact overall scaffold remodeling in a body wall repair setting^7^. The emergence of immunotherapy to treat autoimmune conditions and cancer adds a new dimension to tissue engineering, where immune agonists may be delivered systemically or locally to modulate immunity. For example, we recently showed that subcutaneous co-delivery of the STING pathway agonist cyclic di-AMP (CDA) synergized with ECM scaffold delivery in a therapeutic cancer vaccine while maintaining Type 2 immune signatures like IL-4 expression and STAT6 phosphorylation^8^. Other preclinical studies implanted STING agonist releasing hydrogels following tumor resection surgery, which places a pro-inflammatory agonist in proximity to healing tissue. Although immune modulation is a key determinant of scaffold performance and immunotherapies are becoming prevalent, a detailed relationship between immune agonist and ECM scaffold remodeling has yet to be described.

In the present study (**Figure 1A**), we systematically determine how co-delivery of distinct pro-inflammatory immune agonists with a small intestinal submucosa (SIS) ECM scaffold during VML (a severe traumatic injury) repair affects the acute (1-week) and long-term (8-week) immune response and ultimately scaffold remodeling. We selected 3 immune agonists with relevance to cancer immunotherapy following surgery or with surgical contamination: the STING pathway agonist CDA (a bacterial second messenger and cancer immunotherapy agent), the TLR4 agonist MPLA (Monophosphoryl Lipid A, a derivative of bacterial LPS), and the cytokine GM-CSF (Granulocyte-Macrophage Colony-Stimulating Factor, myeloid cell mediated cancer immunotherapy). Both CDA and MPLA are classically pro-inflammatory though promoting different cytokine profiles and other intrinsic cell effects. GM-CSF mobilizes, recruits, and differentiates myeloid cells including monocytes and neutrophils but does not directly polarize towards pro-inflammatory phenotypes. The local and systemic immune environment was characterized using multispectral flow cytometry and gene expression analysis in muscle, inguinal lymph nodes, and spleen tissues. Histologic analysis was performed for scaffold/tissue remodeling behavior. Additionally, we included control groups receiving soluble immune adjuvants alone to isolate the effects of immunoregulation with trauma alone.

**Figure 1.**
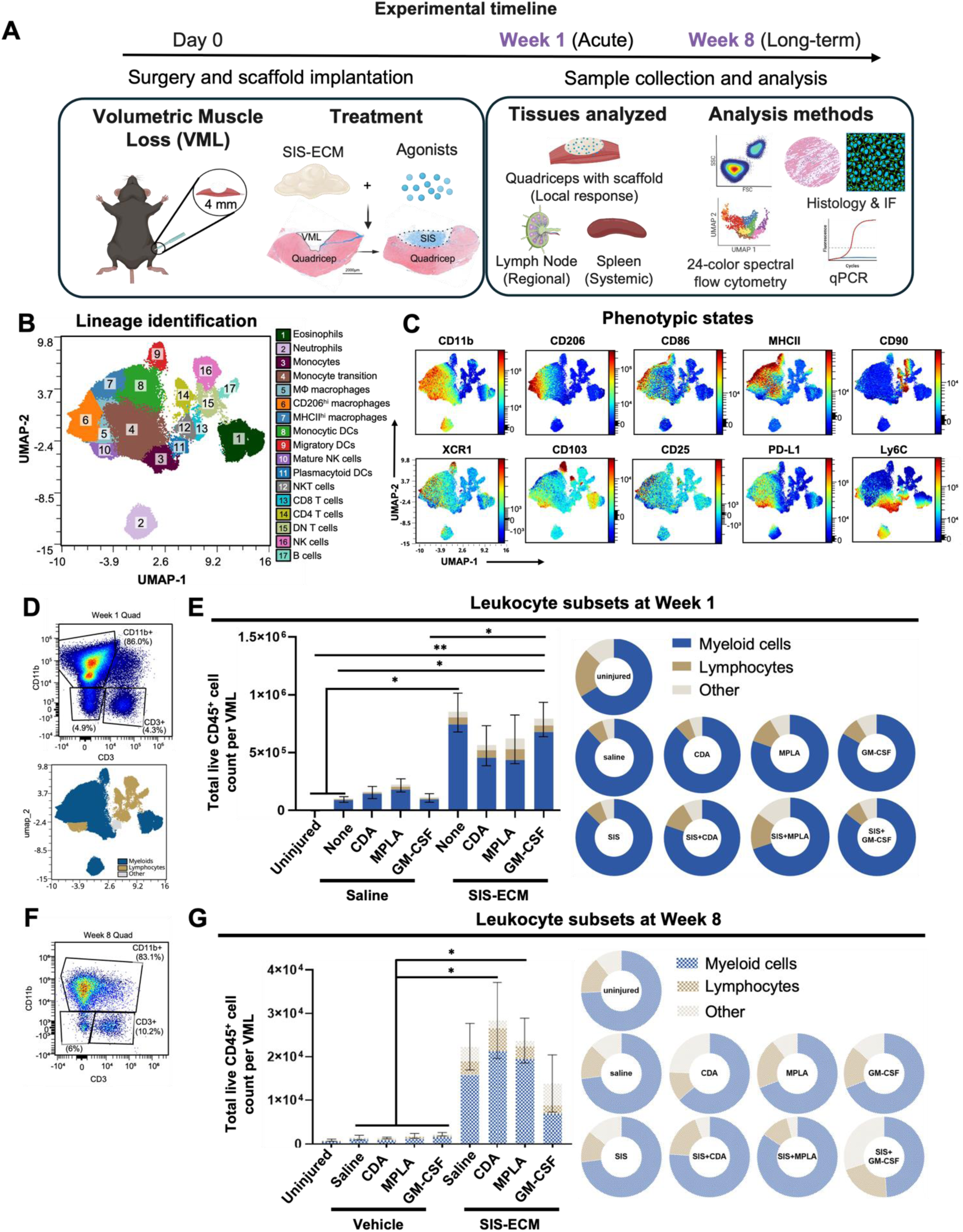
Immunomodulation of the ECM scaffold and VML injury microenvironment with immune agonist co-delivery. **(A)** Schematic of the experimental study design: Female C57Bl/6 mice received volumetric muscle loss (VML) injury to the quadriceps that was repaired with a combination of SIS-ECM scaffold and immune agonist: cyclic di-AMP (CDA), monophosphoryl lipid A (MPLA), or granulocyte-macrophage colony stimulating factor (GM-CSF). The entire muscle, inguinal lymph nodes, and spleen were harvested after 1 week to determine acute inflammatory response and 8 weeks for long-term effects. **(B)** Multispectral flow cytometry was performed for all groups and time points and combined for dimensionality reduction via UMAP and clustering analysis of leukocyte lineages via the FlowSOM algorithm with manual annotation. **(C)** Distribution of markers associated with leukocyte activation and phenotype on UMAP overlay. **(D, F)** The example gating (up) and noted UMAP (below) illustrations of **(D)** week-1 and **(F)** week-8 myeloid cell and lymphocyte populations. **(E,G)** The total number and composition of CD45+ leukocytes invading VML injury and scaffolds across myeloid cell (CD11b+, DCs) and lymphocyte (T cell, B cell, and NK cell) lineages after **(E)** 1 week **(G)** and 8 weeks post-injury. Cell counts analyzed by 1-way ANOVA and posthoc Tukey test: * P < 0.05. Panel A was created in Biorender.

## 2. Results

### 2.1 Type 1 immune agonist co-delivery reduces acute leukocyte density in ECM scaffold treated VML

We applied high-parameter multispectral flow cytometry to reveal the leukocyte lineages and phenotypic states that are modulated when specific immune agonists are locally co-delivered with SIS-ECM scaffolds or saline vehicle controls to treat severe trauma. A 24-color panel identified 17 unique leukocyte clusters within VML injured quadriceps tissue via dimensionality reduction and supervised clustering (**Figure 1B**), which included significant heterogeneity in activation state (**Figure 1C, SFigure 1A**). This data-driven population identification approach defined populations, including complex myeloid cell phenotypes like macrophages (**SFigure 1B,C**), that largely agreed with manually gated populations defined *a priori* (gating strategy provided in **SFigure 2**). Manual annotation was required to separate relatively low feature subsets such as B cells (cluster 17) from double negative (DN) T cells (cluster 15) using expression of B220. The cluster map was primarily segregated into regions enriched for monocyte-macrophage-dendritic cell states, lymphocytes, and granulocytes (neutrophils and eosinophils). As expected, antigen presentation machinery MHCII was found in subsets of macrophage, dendritic cell (DC), and B cell populations. Likewise, reparative CD206^+^ macrophages that are characteristic of the response to ECM biomaterials were identified as well as hybrid macrophages co-expressing the classical M1 and M2 markers CD86 and CD206, respectively.

**Figure 2.**
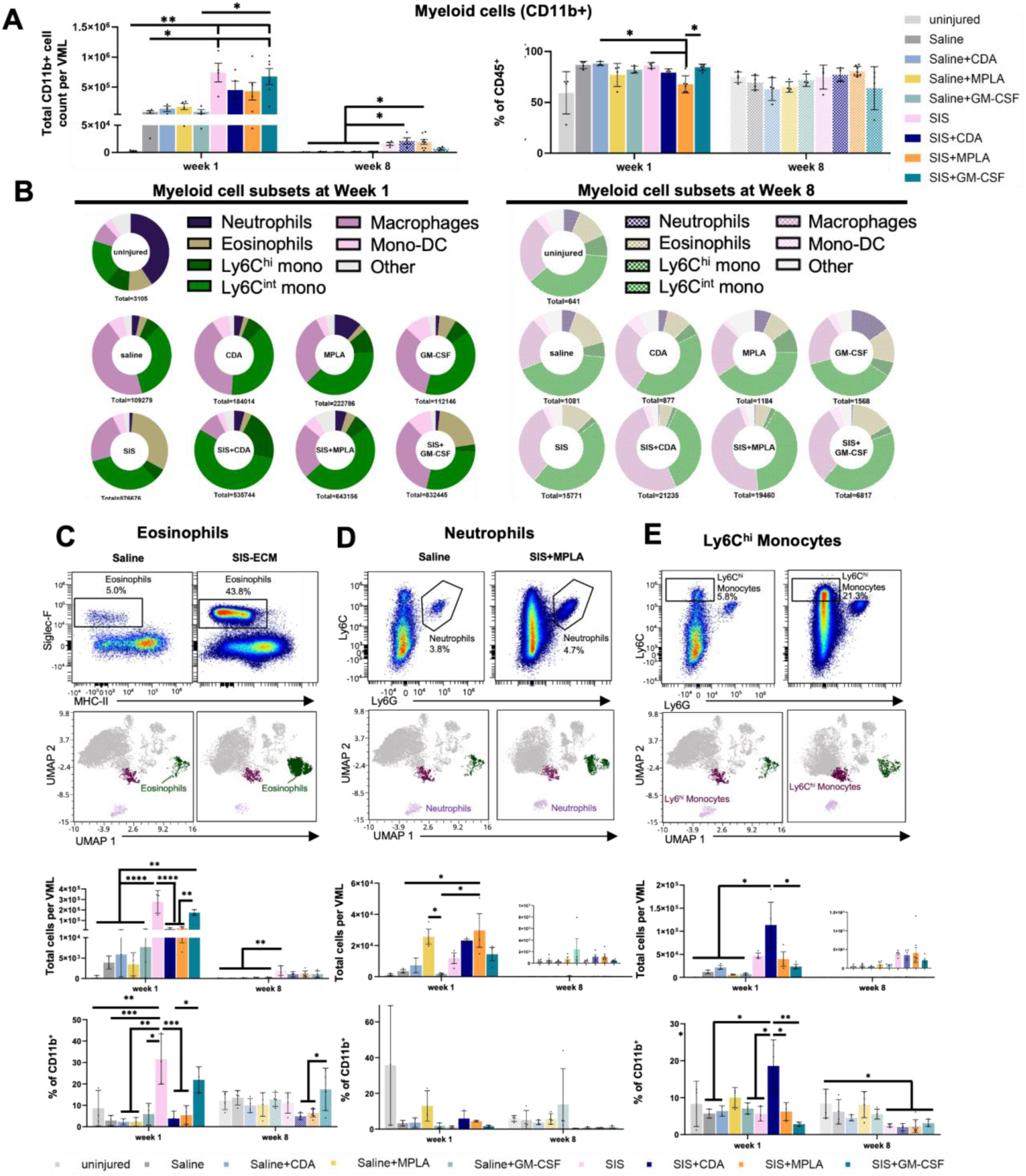
ECM scaffold induced changes in myeloid cell composition, polymorphonuclear cells and monocytes. **(A)** Total CD11b^+^ myeloid cell as cell count (left) and frequency of leucocytes (right) at week 1 and 8. **(B)** Myeloid cell composition changes at week 1 (left) and week 8 (right) with different treatments. **(C)** eosinophils, **(D)** neutrophils, and **(E)** Ly6C^hi^ monocytes flow cytometry gating, UMAP population, cell count and percentage comparison among treatments. Cell counts and percentage comparison analyzed by 1-way ANOVA and posthoc Tukey test: * P < 0.05.

UMAP and cluster analysis revealed additional insights into the VML and scaffold immune microenvironment (**Figure 1C**) that were not expected from our prior knowledge-driven manual gating approach. Monocytes (cluster 3) exhibited a qualitative spectrum of differentiation within the VML repair site, transitioning Ly6C^+^ monocytes (Cluster 4) to macrophage (clusters 5, 6, 7) and dendritic cell (cluster 8, CD11b+ monocytic DC) trajectories. Unexpectedly, there was a myeloid cell subset resembling mature NK cells (NK1.1+ CD11b+) clustered closely with M0 macrophages that were among the highest expressors of the inhibitory ligand PD-L1. The tissue homing integrin CD103 was expressed by their canonical cell types in migratory DCs and T cells but also discovered on eosinophils and mature NK cells. Likewise, the antigen cross-presenting marker XCR1 was found on M2 macs and not only dendritic cells. CD25, the IL2 receptor subunit found in regulatory T cells and activated T cells, was found distributed among antigen presenting macrophages expressing MHCII.

We subsequently quantified immune populations (both the traditional and unexpected phenotypes) using manual gating to determine the ECM scaffold and immune agonist specific drivers. The absolute cell numbers reflect the intensity of inflammation and relative abundances (as % within a lineage) determines the balance of these immune programs within the VML. Mature leukocytes were defined as viable CD45^hi^ cells; though CD45^mid^ cells and CD45^−^ stromal cells were also observed following injury but not in uninjured muscle. Consistent with previous studies, ECM scaffold implantation resulted in an 8-fold increase in the total number of leukocytes recruited to the scaffold after 1 week (1.0×10^5^ vs 8.5×10^5^ CD45^+^ cells/VML) in the absence of exogenous immune agonist stimulation. The addition of any of the tested immune agonists alone in a saline vehicle did not substantially increase leukocyte density in the VML site (**Figure 1D**). However, an interaction effect occurred with ECM scaffolds. GM-CSF delivered with SIS-ECM increased immune recruitment compared to GM-CSF alone (7.9×10^5^ vs. 1.1×10^5^ CD45^+^ cells/VML, respectively). Unexpectedly, incorporating pro-inflammatory immune adjuvants slightly blunted immune recruitment, with MPLA (6.2 ×10^5^) and CDA (5.6 ×10^5^ CD45^+^ cells/VML).

Myeloid lineage cells (**Figure 1E**) were the most abundant acute (1-week) leukocyte population (87%) in both saline and SIS-ECM treated VMLs, followed by lymphocytes (7%) and other immune cells (5%). MPLA was the most potent agonist in reshaping the VML immune composition, reducing myeloid cell proportion (80% with saline and 69% with SIS) while expanding lymphocytes (11% with saline and 15% with SIS). CDA moderately affected the SIS immune environment relative to MPLA with a reduction in myeloid cells (80%) and increased lymphocytes (11%); however, CDA did not affect composition when delivered alone in saline. GM-CSF did not affect these immune proportions.

While pro-inflammatory adjuvants CDA and MPLA reduced acute inflammation within SIS ECM scaffold treated injury, they promoted long term immune persistence. By 8 weeks (**Figure 1F,G**), VML trauma inflammation had largely resolved to uninjured levels in all saline groups regardless of adjuvant delivery (0.8-2.2×10^3^ CD45^+^ cells/VML). ECM scaffold implantation leukocyte recruitment was slightly elevated (2.2×10^4^ CD45^+^ cells/VML) and approximately 20-fold fewer cells than at 1 week. CDA and MPLA potentiated SIS-ECM scaffold inflammation, inducing elevated cell densities at 2.8 and 2.3×10^4^ CD45^+^ cells/VML, respectively. GM-CSF in contrast was not as substantially elevated 1.3×10^4^ CD45^+^ cells/VML. Myeloid cells remained the greatest leukocyte subpopulation, followed by similar lymphocyte and other immune cell proportions. These results show that SIS-ECM scaffold implantation facilitates a response to immune agonist in trauma, as immune agonist delivery alone resolves by 8 weeks but potentiates the SIS ECM response.

### 2.2 ECM scaffold polymorphonuclear cells and monocytes are differentially modulated by co-delivery with specific immune agonists

Myeloid lineage cells of innate immunity are crucial to both the early and late immune response to biomaterials, with polymorphonuclear (PMN) cells such as neutrophils and eosinophils as the earliest responders transitioning to monocytes then to macrophages and dendritic cells. We found that each immune agonist generated a unique myeloid recruitment signature within SIS-ECM scaffolds after 1-week. Total CD11b^+^ myeloid cell density (**Figure 2A**) correlated with total leukocytes (**Figure 1**), in which SIS-ECM with and without GM-CSF enhanced myeloid cell recruitment by 7.6- and 8-folds relative to injury alone. CDA and MPLA blunted myeloid recruitment early but persisted by 8 weeks. Myeloid cells composed a similar proportion of total leukocytes (∼83%) in all groups except with MPLA and SIS-ECM which decreased to 73%.

We closely examined how agonist delivery altered scaffold PMN and monocyte-macrophage-dendritic cell phenotypes in VML. Based on prior UMAP analysis, there was a spectrum of differentiation from classical monocytes (Ly6C^hi^ mono) to intermediate states (Ly6C^int^ mono) to fully matured monocyte derived macrophages and dendritic cells (Mono-DC). The proportions of these populations varied by scaffold and immune agonist condition, beginning with early responders in eosinophils, neutrophils, and Ly6C^hi^ monocytes (**Figure 2B-F**). Eosinophils (Siglec-F^+^ MHCII^lo^), a hallmark of the Type 2 immune response to ECM scaffolds derived from animal tissues^9,10^, showed one of the greatest differences between saline and SIS-ECM alone treatments. Rising from 3,480 cells (2.6% of CD45⁺ cells) in saline to 277,154 cells (27.7%) with SIS-ECM at week 1, a trend that persisted through week 8 (**Figure 2B,C**).

The addition of GM-CSF maintained eosinophil bias in SIS-ECM, however the pro-inflammatory adjuvants CDA and MPLA both substantially reduced eosinophil recruitment by approximately 17- and 10-fold. Immune agonists alone did not significantly affect eosinophil presence. Neutrophils are known to be involved in the hyper-acute phase of injury and ECM inflammation (1-3 days^8^) but is largely gone after 1 week in the absence of additional stimulus and is an indicator of a Type 1 response. Indeed, MPLA increased neutrophil mobilization alone and with ECM scaffolds and was the only agonist to modulate any myeloid cell type when delivered to the VML injury environment alone (**Fig 2E**). Neutrophil recruitment was similar between ECM scaffold and saline delivery indicating that SIS-ECM does not synergistically participate in this orthogonal response. Mean neutrophil density was at background uninjured levels by 8 weeks in all groups. Ly6C^hi^ monocytes following PMN recruitment are precursors to CD11b^+^ macrophages and DCs. Interestingly, we found that CDA markedly increased monocyte presence, 1.1×10^5^ cells versus 4.7×10^4^ with SIS-ECM alone and raised their proportion from 5.4% to 21.2%. In summary, each immune agonist showed a distinct myeloid signature bias in SIS-ECM: CDA enhanced monocytes and decreased eosinophils, MPLA enhanced neutrophils and decreased eosinophils, and GM-CSF promoted accumulation of macrophages and DCs while maintaining eosinophil presence. Similar to overall leukocyte recruitment, immune agonist activity is primarily detected within the SIS-ECM environment even during the early stages of healing, with the exception of MPLA recruiting neutrophils in the absence of scaffold.

### 2.3 CDA switches macrophage polarization and antigen presenting cell activity in the SIS-ECM microenvironment

The earliest phase of inflammation following injury and biomaterial implantation transitions from PMNs to monocyte differentiation into macrophages and dendritic cells. Each of these innate immune cells are phenotypically and functionally plastic depending on stimuli received from their environment via receptors to the agonists tested here. Both clustering analysis and manual gating showed that there was a spectrum of monocyte differentiation; we found a continuum of Ly6C downregulation suggesting active differentiation into Ly6C^−^ macrophages (as determined via FMO controls). These transitional Ly6C^int^ and mature Ly6C^−^ populations gained expression of the macrophage marker F4/80 during macrophage differentiation, the dendritic cell integrin CD11c, or both (**Figure 3A**). We found that mature macrophages (Ly6C^−^) showed the greatest co-expression of CD11c and F4/80, which has been described previously though their functional significance is unknown.

**Figure 3.**
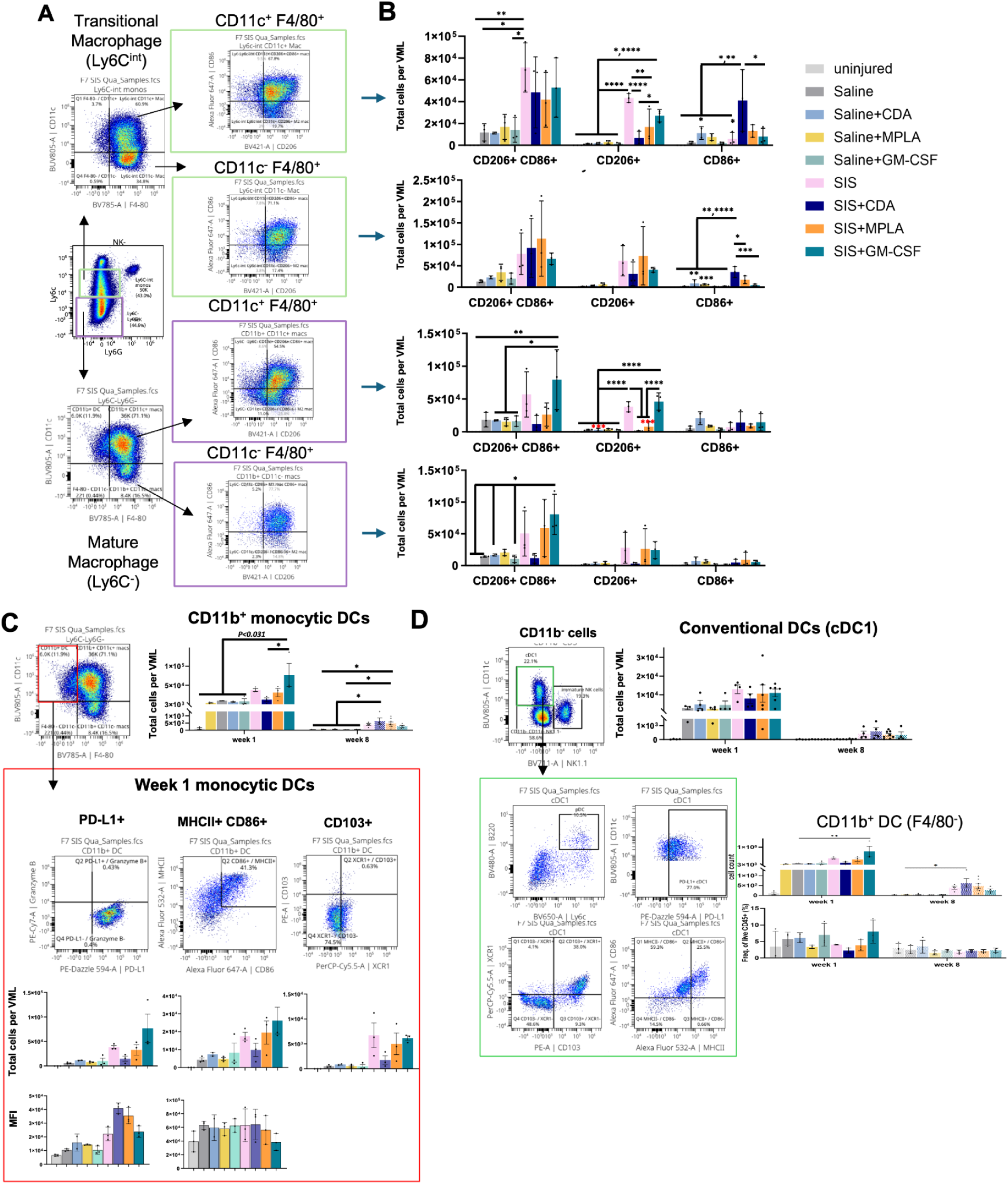
ECM scaffold induced changes in macrophages and dendritic cells. **(A)** Macrophage subsets gating. Transitional stage macrophages were gated by Ly6C-intermediate and F4/80+ expression. Then macrophages were subgated based on CD11c co-expression. **(B)** Macrophage subsets CD86, CD206 or hybrid cell counts comparison. **(C)** Week-1 monocytes derived-dendritic cells and **(D)** CD11b-conventional dendritic cells gating, cell count and activation states comparison. Cell counts and percentage comparison analyzed by 1-way ANOVA and posthoc Tukey test: * P < 0.05.

We then determined how immune agonist delivery modulated macrophage phenotype with and without SIS-ECM scaffold mediated repair across the spectrum of transitional to mature macrophages. Macrophage phenotype is complex, however surface markers induced by specific stimuli under ideal in vitro conditions are a useful categorization framework. M2 macrophages are enriched with exposure to the Type 2 cytokine IL-4 and M1 macrophages in response to lipopolysaccharide (of which MPLA is a subunit) and the Type 1 cytokine IFNγ. ECM scaffold recruited macrophages are heterogeneous but overall biased towards M2-like macrophages and hybrid phenotypes that express both CD206 and CD86^9,11^. Our results with SIS-ECM alone were consistent with that finding; as the majority of these cells were CD206^+^ or hybrids in each subset after 1 week compared to saline treatment with or without agonist (**Figure 3B**). Agonist delivery did not have a strong effect on hybrid Ly6C^int^ transitional macrophage polarization in the SIS-ECM microenvironment. CDA greatly reduced CD206^+^ expressing cells and increased CD86^+^ expression, in both CD11c^+/-^ subsets. MPLA had a smaller magnitude decrease in CD206 but no effect on CD86 in CD11c^+^ macrophages, and GM-CSF was generally similar to SIS-ECM. In mature Ly6C^−^ CD11c^+^ macrophages, CD206 was substantially reduced with both CDA and MPLA co-delivery. However, CD86 expression was not affected by CDA suggesting it has less of a role in regulating mature macrophages. Macrophage expression patterns with GM-CSF matched SIS-ECM alone. We corroborated agonist effects by examining the markers MHCII (major histocompatibility complex class II) and PD-L1 (programmed cell death protein ligand 1), which are crucial for antigen presentation during adaptive immune response activation and as a negative feedback regulator of adaptive immunity, respectively. The majority of macrophage (> 97%) expressed PD-L1 indicating activation (SFig 1). In contrast, MHCII expression mirrored hybrid macrophage abundance, with a strong increase in with ECM scaffolds across subsets, including CDA-induced decreases in MHCII in mature macrophages. After 8 weeks, macrophage numbers declined to fewer than 2,000 per VML, with no substantial regulation per group; the majority of which being hybrid or CD206^+^ subsets. In summary, agonist alone didn’t have much of an effect on macrophage polarization alone though its effect on ECM was generally to promote Type I immunity with CDA and MPLA, and to preserve Type II immunity with GM-CSF.

Dendritic cells are highly specialized innate immune cells that internalize and present antigenic material and provide stimulatory signals to T cells. Monocyte derived (CD11b^+^ F4/80^−^) were less numerous than macrophages in all groups, though there was a significant increase with SIS-ECM and SIS-ECM+GM-CSF. PD-L1 expression dynamics were different from macrophages, with increased expression in CDA and to a lesser extent MPLA (**Figure 3C**). Mature DCs are defined as co-expression of both MHCII and CD86 as necessary for T cell priming, where there is a slight reduction in CDA numbers, but not expression intensity. Lastly, CD103 indicates a migratory phenotype, and overall, very few of these cells were observed.

In addition to DCs derived from monocytes mobilized from the blood, tissue resident conventional DCs (cDCs) respond to injury and biomaterial implantation^12^.

### 2.4 The adaptive immune response to SIS-ECM scaffolds is potentiated by immune agonist delivery

Adaptive immune activation is a highly coordinated series of events downstream of early innate activation that was recently shown to be a critical regulator of the host response to biomaterials. Immune stimuli polarize the myeloid cells that prime lymphocytes such as T cells or can interact with these cells directly via many of the same pattern receptors. Previous studies have shown that T cells, primarily CD4^+^ helper T cells respond to ECM scaffold implantation, with an order of magnitude fewer CD8^+^ cytotoxic T cells or B cells^9,13^. Our results showed that SIS-ECM and immune agonist altered the lymphocyte profile within volumetric trauma, with a greater proportion of T cells with either ECM or agonist after 1 week, that returned to baseline after 8 weeks (**Figure 4A**). We then examined the CD4 T helper cell, CD8 cytotoxic T cell, and double negative (DN) subsets (**Figure 4B**). As expected, CD4 T helper cells were the most abundant, comprising greater than 80% of T cells in SIS-ECM, followed by DN and CD8 T cells (**Figure 4C,D**). CDA increased the proportion of CD8 cytotoxic T cells when delivered via saline or with SIS-ECM. Unexpectedly, DNT cells were increased with the pro-inflammatory adjuvants in CDA and MPLA. DNT cells play a regulatory role in maintaining tissue homeostasis and can be expanded in sites of injury^14^, though their presence in ECM-scaffolds have yet to be described. Mature NK cells (CD11b^+^) and immature NK cells (CD11b^−^) were enriched in ECM scaffolds but not regulated by adjuvant. Though SIS-ECM effectively recruited CD4 cells, there were few effects of adjuvant on these broad categories. We then examined T cell activation state and canonical Th2 and Th1 cytokine expression, *Il4* and *Ifng*, respectively. In agreement with prior studies, *Il4* but not *Ifng* expression is greatly enhanced with SIS-ECM implantation in soft tissue trauma (**Figure 4E**). The agonists CDA and GM-CSF decreased Ill4 expression relative to SIS alone, but it remained 10-fold greater than untreated saline. Surprisingly, MPLA did not influence Il4 expression, nor did any agonist alone delivered with saline. Likewise, CD8 T cell activation state did not change via CD25 and granzyme B, though we found that CDA drove CD4 T cell activation (**Figure 4F**) without a concomitant increase in regulatory T cell abundance (T_reg_).

**Figure 4.**
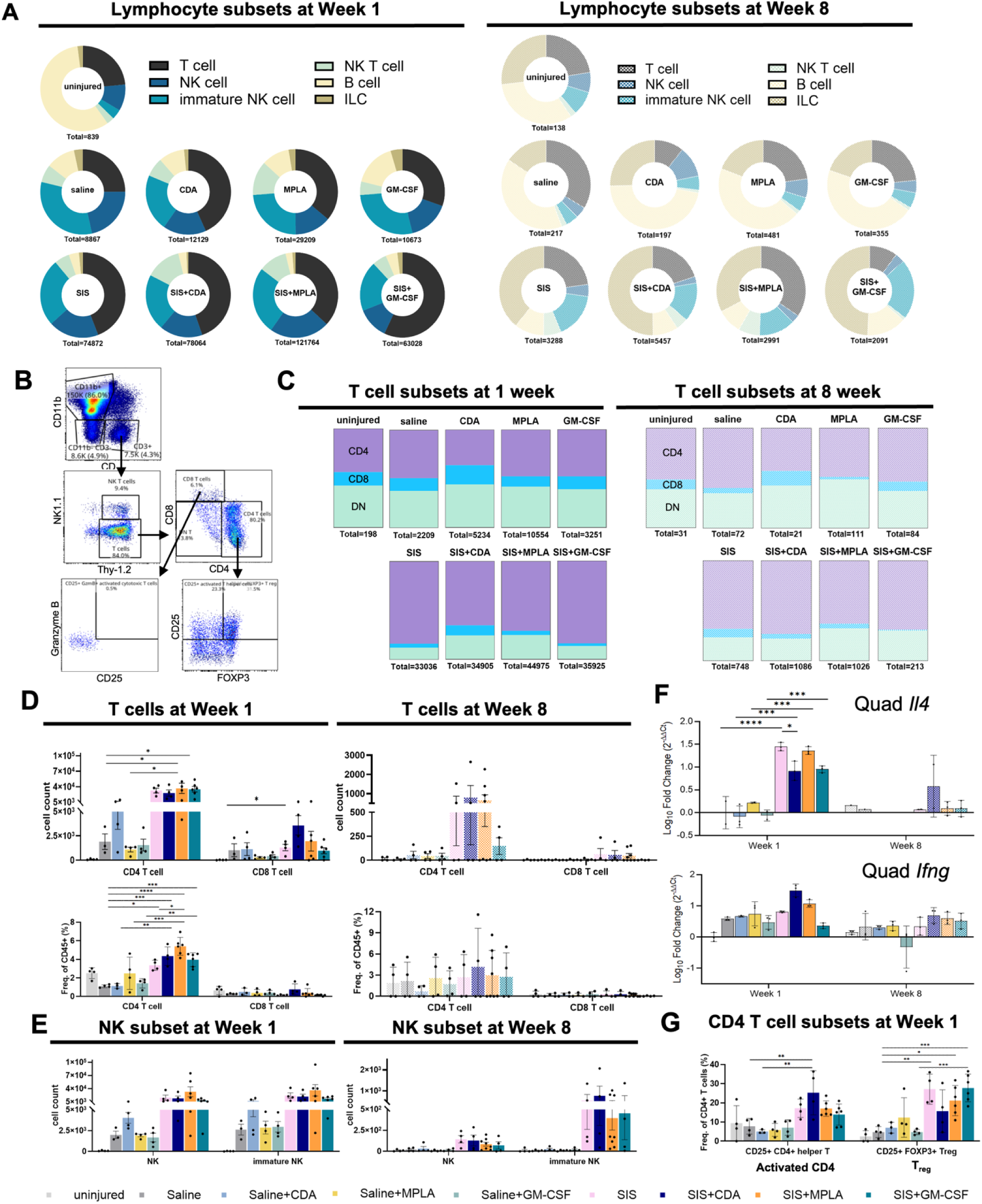
ECM scaffold induced changes in lymphocytes and cytokine expression. *(A)* Lymphocytes subset compositional comparison among treatment groups at week 1 and 8. **(B)** T cell lineage and activation markers gating. **(C)** T cell subset comparison among treatment groups at week 1 and 8. **(D)** T cell and **(E)** NK cell subset compositional comparison among treatment groups at week 1 and 8. **(F)** Quadricep cytokines expression at week 1 and 8. **(G)** CD4 T cell activation state comparison at week 1. Cell counts and percentage comparison analyzed by 1-way ANOVA and posthoc Tukey test: * P < 0.05.

### 2.5 Immune agonist CDA enhances CD86 expression in adjacent muscle while GM-CSF accelerates SIS-ECM degradation

We performed histologic analyses to determine the spatial distribution of the canonical polarization markers CD206 and CD86 and subsequent effect on scaffold remodeling. After 1-week, untreated VML injuries demonstrated localized inflammation, with macrophages expressing CD86 in adjacent muscle tissue (**Figure 5A**). SIS implantation enhanced total cellularity and the presence of CD206+ cells. The adjuvants GM-CSF and MPLA had nominal effects on inflammation. However, CDA induced significant changes to nearby tissue. There was extensive muscle fiber death, interspersed with dense accumulation of CD86^+^ cells (both F480^+/-^ cells). This accumulation occurred with saline and SIS-ECM delivery but was most pronounced with SIS-ECM.

**Figure 5.**
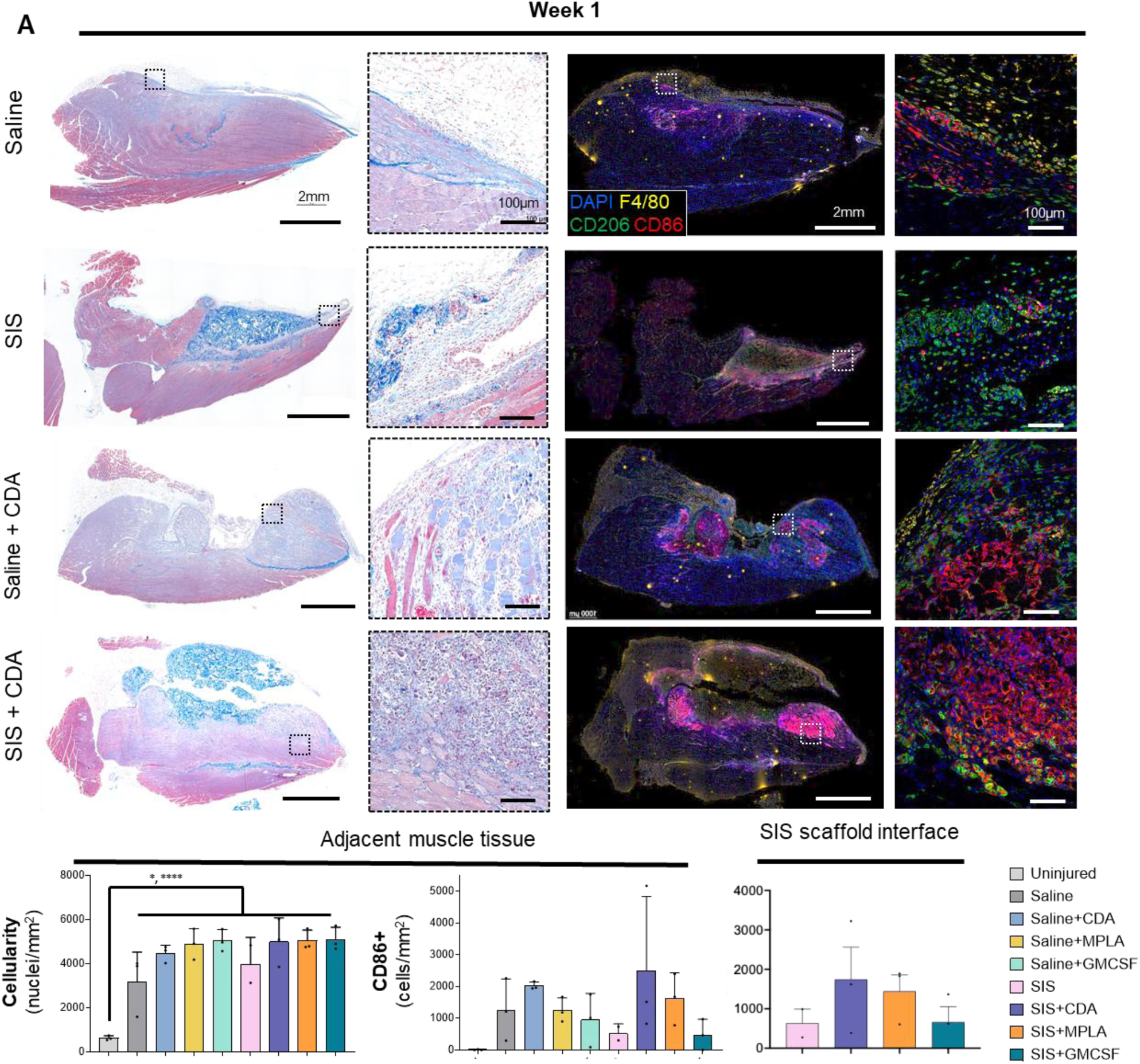
Histologic and spatiotemporal immune profiling of VML injury post week 1 of ECM scaffold implantation. **(A)** Masson’s Trichrome-stained images (Left panel) and multiplex immunofluorescent images (Right panel) showing the morphology of VML injury microenvironment with immune agonist co-deliver (CDA, MPLA, or GM-CSF, **S**Figure 5) with SIS-ECM scaffold or saline post 1 week of injury and implantation in C57Bl/6 mice quadriceps. Boxes highlight immune infiltrates at the at 20X objective. (**B**) Total cell density (N=3, mean ± SD) and CD86+ myeloid cell density in the adjacent muscle tissue of injured quadriceps. (**C**) CD86+ myeloid cell density in the SIS scaffold interface of injured quadriceps (N=3, mean ± SD). *p < 0.05, ****p < 0.0001 one-way ANOVA with Tukey’s multiple comparisons test.

We then examined long-term effects of this immunomodulation (**Figure 6**). All saline repair groups with or without adjuvant were histologically identical, regardless of adjuvant type. SIS-ECM had partially degraded and remodeled into adipose and loose connective tissue, with occasional muscle fiber formation around the edges. The adjuvant GM-CSF promoted the greatest remodeling effect, encouraging the most degradation and new tissue deposition.

**Figure 6.**
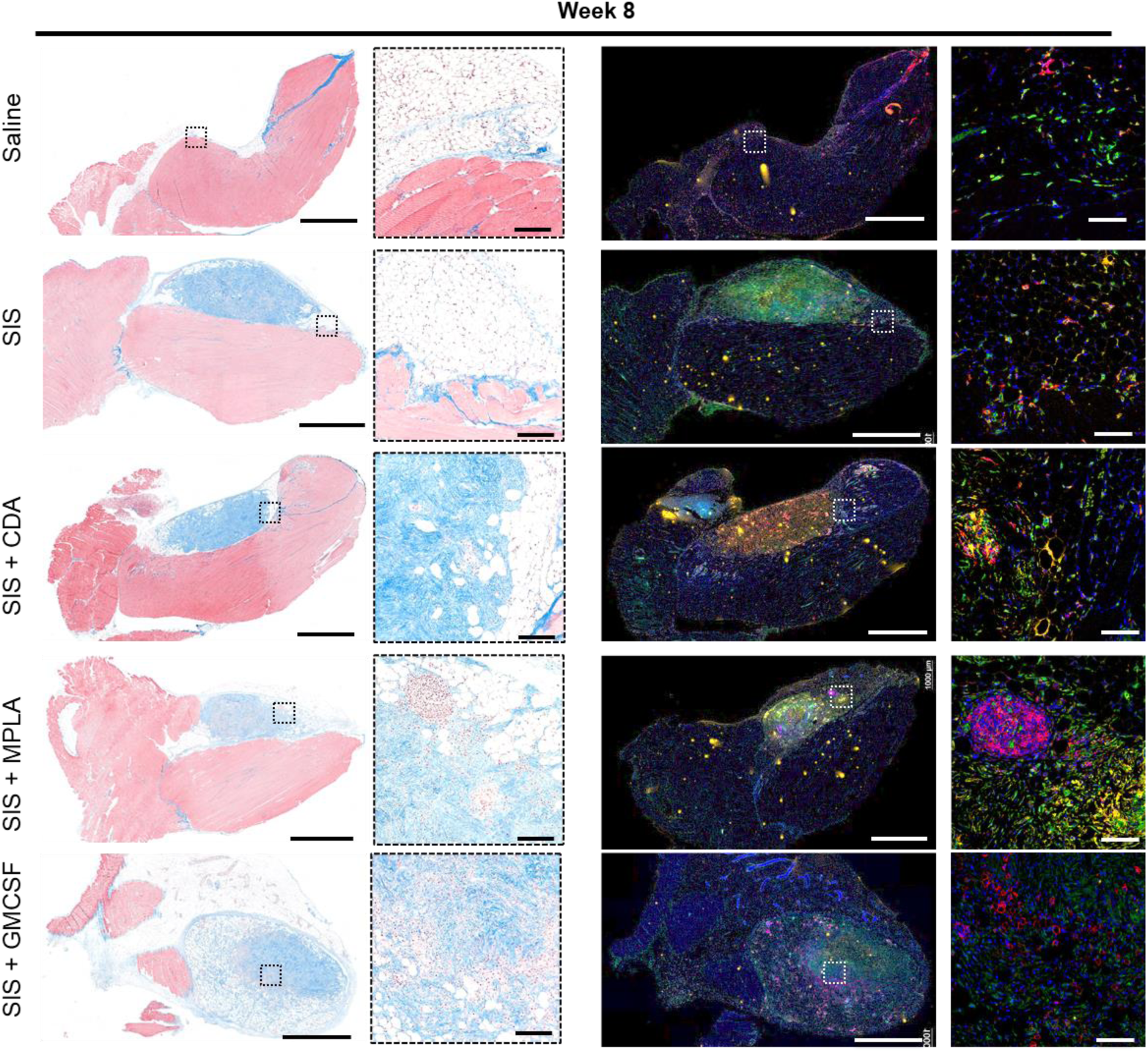
Histologic and spatiotemporal immune profiling of VML injury post week 8 of ECM scaffold implantation. Masson’s Trichrome-stained images (Left panel) and multiplex immunofluorescent images (Right panel) showing the morphology of VML injury microenvironment with immune agonist co-deliver (CDA, MPLA, or GM-CSF) with SIS-ECM scaffold (saline controls in **S**Figure 6) post 8 weeks of injury and implantation in C57Bl/6 mice quadriceps. Boxes highlight immune infiltrates at the at 20X objective.

## Discussion

In the present study we determined how activating specific inflammatory pathways with infection mimicking immune agonists used in vaccine immunotherapy reshapes the ECM biomaterial scaffold microenvironment during healing and remodeling after musculoskeletal trauma. High parameter multispectral flow cytometry revealed the agonist specific immune signatures that develop during the acute inflammatory phase of injury and scaffold implantation. Though all agonists have inflammation inducing actions, their effect on the ECM microenvironment was unique. The TLR4 agonist MPLA increased neutrophil content, reduced eosinophil infiltration, and promoted increased CD4^+^ T cell density; the STING agonist CDA promoted monocyte recruitment and decreased eosinophils while shifting CD206^+^ macrophages to CD86^+^ macrophages. GM-CSF did not alter Type 2 immune features such as eosinophils and CD206^+^ macrophages but promoted the increased density of monocyte-derived DCs and lymphocytes. Some Type 2 features remained such as high *Il4* expression that was elevated in all ECM groups. Unexpectedly, these same immune agonists induced few changes to the immune environment when delivered in a soluble form directly into the wound without an immunomodulatory ECM scaffold. This suggests a highly transient effect of these agonists alone and an interaction effect between the pro-reparative ECM immune response. Ultimately, these agonists affected tissue remodeling and immune infiltration in nearby injured tissues. CDA promoted high CD86 dense areas in nearby muscle after 1 week, but little effect on remodeling after 8 weeks, whereas GM-CSF strongly promoted scaffold degradation and development of loose connective tissues and adipose.

We were motivated to evaluate immune agonists in muscle healing because of the developing intersection of surgery and ECM scaffold repair with immunology and immunotherapy. Infection remains an ever-present risk factor in traumatic injury. Prior literature has shown that ECM scaffolds can be advantageous in wounds with high contamination risk. Antimicrobial features for ECM biomaterials are their degradability (preventing durable biofilm formation) and direct microbial inhibition from ECM proteolytic degradation products released during remodeling. Deliberate infection studies showed that microbial growth inhibited, but detailed immune characterization and its effect on long-term remodeling was not fully elucidated. There is conflicting evidence in hard tissues of whether the immune response to infection accelerates or impairs quality osteogenesis, and we sought to determine if targeted immune modulation could be beneficial to ECM mediated repair.

Likewise, immunotherapy is a rapidly growing therapeutic class, which intersects with tumor resection surgery and tissue repair. Our previous work showed that the inflammation stimulated by an ECM scaffold was synergistic with a therapeutic cancer vaccine immunotherapy, however, this was in the absence of traumatic injury that would occur after tumor surgery. MPLA promoted neutrophil recruitment and CDA showed a reduction in macrophage density at 1 day and 1 week post-implantation, which would be consistent with the monocyte dominant response to CDA here. *Il4* expression was high in that study and increased with agonist. More prominent regulation was observed here, suggesting that the wound healing response from traumatic injury is a major contributor to adjuvant response with ECM scaffolds. We unexpectedly found that the full combination of ECM scaffold and injury were necessary to potentiate the effect of agonist.

Ultimately, we were interested in connecting acute inflammatory regulation of these agonists on local tissue repair. There were no adverse events or evidence of systemic inflammation observed in this study. Animals were fully ambulatory throughout the study and systemic inflammation hallmarks such as splenomegaly were absent which confirms a localized immune response. However, we did observe that the increased CD86 expression observed with CDA delivery was localized to dense areas in adjacent tissues where muscle degeneration is occurring. We did not observe loss of muscle mass or fibrosis after 8 weeks, which would support STING pathway induced muscle damage though functional assays are required to confirm. However, tissues treated with CDA alone, as with all other agonists, were indistinguishable from untreated muscle injury which also supports lack of durable effects on muscle healing. Notably, these dense lakes of CD86, a co-stimulatory molecule in T cell activation, has immunological implications. There may be heightened immune surveillance against pathogenic agents such as microbes or neoplastic cells in the local region that is transient, without chronic inflammation. Conversely, autoimmunity is a risk with chronically elevated adaptive immune stimulatory conditions should self-tolerance be broken. We did not observe any such evidence of autoimmunity here, even after 8 weeks which is sufficient to induce autoimmune conditions such as arthritis in mice. This may be due to the immunoregulatory nature of ECM or lack of autoantigen release under these conditions.

We found that specific immune agonist delivery affected the long-term remodeling of ECM scaffolds. Our initial hypothesis was that increased inflammation induced by these agonists would accelerate degradation rate, with agonist type influencing the type of tissue. However, only GM-CSF and MPLA qualitatively affected scaffold degradation, with CDA delivery with SiS-ECM having a nominal effect. This heightened remodeling with GM-CSF correlated with increased mature macrophage and DC density, which was not induced by the other adjuvants. Though these are infection-mimicking compounds did not promote fibrotic scar tissue formation after 8 weeks, which is a consequence of chronic inflammation or the foreign body response to nondegradable biomaterials. Agonists did not promote this fibrotic program. Alternatively, there was increased connective tissue and intermittent adipose tissue formation, which is one of the reported outcomes of fibro-adipogenic progenitors. We did not observe gross changes in muscle architecture, but additional analysis of myogenesis is necessary to fully characterize this outcome. These results may have therapeutic implications for the type and location of immunotherapy following surgical trauma.

## 3. Experimental Section

### 3.1. Preparation of Decellularized Small Intestinal Submucosa (SIS) Extracellular Matrix (ECM) Particles

Porcine small intestine was obtained from Tissue Source LLC (Zionsville, IN) from market-weight pigs that were documented as pathogen-free (porcine reproductive and respiratory syndrome, porcine epidemic diarrhea virus, porcine delta coronavirus, transmissible gastroenteritis) and complied with ISO 13485 and stored frozen at -30°C. Small intestines were thawed at 4°C overnight and flushed of their contents, cut open along its length, and then mechanically delaminated to remove the muscularis and mucosal layers. The remaining submucosa was cut into 6-inch pieces and decellularized using 4% alcohol/0.1% peracetic acid (v/v, Sigma-Aldrich), washed thoroughly with 1X PBS and Type 1 water, and then lyophilized and comminuted into an injectable particulate form via cryogenic milling and passing through a 425 µm pore sieve. SIS particles were terminally sterilized via 2×106 rad gamma irradiation on dry ice. Particles were tested as negative for common murine viral and bacterial pathogens before use in animal experiments.

### 3.2. Volumetric Muscle Loss (VML) Model

Ethical approval for the animal experiments was provided by the Institutional Animal Care and Use Committee at NCI Frederick (ASP No. 23-063). C57BL-6J female mice were sourced from The Jackson Laboratory and kept in the NCI Frederick Laboratory Animal Sciences Program in specific pathogen-free conditions and under 12-hr light/dark cycles for at least 1 week of equilibration prior to experimental procedures at 8 weeks old. Mice were euthanized via asphyxiation with carbon dioxide and cervical dislocation.

Bilateral volumetric muscle loss injuries were inflicted to evaluate the host remodeling response to implant. Mice were anesthetized using 1.5-2.5% isoflurane in oxygen for surgical procedures in a biosafety cabinet and provided subcutaneous Buprenorphine SR at 1 mg/kg for analgesia. The surgical site was scrubbed with chlorhexidine and 70% ethanol aseptically 3 times, followed by a volumetric muscle loss injury in the quadriceps as previously described ^15^ . Briefly, a 1 cm unilateral longitudinal incision was made in skin and muscle fascia to expose the underlying quadricep muscle, after which a 4 mm long by 3 mm deep section was removed from the center of muscle body to create a critical sized muscle deficit. Defects were either filled with a saline control, adjuvant-saline suspension, ECM particulate material hydrated with saline, or ECM particulate material hydrated with an adjuvant-saline solution. 20 µg CDA, 10 µg MPLA, or 1 µg GM-CSF was prepared in 50 µL of saline, and correspondingly combined with 10 mg of ECM to fill the muscle deficit. After implantation, the overlying skin was closed using surgical staples and 4-0 Vicryl sutures, and a topical 0.25% Bupivacaine solution was applied to the incision site. The procedure was repeated on the other leg. Mice were monitored daily for the first 3 days following surgery, then weekly after. Motivation for mouse euthanasia included wound dehiscence or other complications that cause distress or affect ambulation.

The entire quadriceps muscle with scaffold, spleen and draining lymph nodes (N=3-4) were harvested in RPMI media for flow cytometry. Another set of quadriceps (N=3) were harvested by reflecting overlying skin and the entire leg was fixed in 10% neutral buffered formalin to preserve tissue morphology for histology analysis.

### 3.3. Histologic analysis and multiplex immunofluorescence staining

Formalin-fixed tissues were dehydrated with a graded series of ethanol and xylene for paraffin embedding, sectioned (5 um), and stained for Masson Trichrome as per standard protocols (Histoserv, Inc.). Masson Trichrome-stained quadriceps were imaged using a Zeiss AxioObserver at 20X objective (high resolution) or 10X objective tiled images.

Immunofluorescence staining and imaging were performed to characterize immune cell infiltration within SIS-ECM scaffolds in combination with pro-inflammatory immune agonists. Sections were deparaffinized in xylene followed by rehydration in decreasing concentrations of ethanol and then in Type I water. Rehydrated sections were post-fixed with neutral buffered formalin for 15 min followed by a wash with Type I water. Antigen retrieval of tissue sections was performed in citrate buffer (pH 6.0) for 20 min at 95-98°C using a steamer and cooled at RT for 20 minutes followed by washing with Type I water. Endogenous peroxidases were quenched by incubating the slides in 3% H2O2 in PBS for 15 minutes. Slides were washed with Type I water, and a border was created around the section using a PAP pen. Unreacted aldehydes were quenched with 2.24% (0.3M) Glycine (w/v) in TBS-T buffer (Tris Buffered Saline with 0.05% Tween-20) for 5 minutes followed by blocking using 10% BSA in TBS-T for 30 min at room temperature.

Sections were sequentially stained via tyramide signal amplification. In brief, each round of staining consisted of incubation with primary the antibody diluted in blocking buffer, 3 washes in TBS-T, incubation with species-appropriate HRP-Polymer conjugated anti-IgG secondary for 15 minutes at room temperature (RT), 3 washes in TBS-T, incubation with Opal dye (Akoya) diluted in Opal amplification diluent (Akoya), and 2 water washes. Antibody stripping was performed between rounds using citrate buffer (pH 6.0) for 20 min at 95-98°C. After the final round, slides were counterstained with DAPI (1µg/ml) in PBS for 5 min followed by water washes and coverslipping with fluorescent antifade mounting reagent (DAKO, Agilent).

For macrophage phenotype staining, primary antibody labeling conditions and Opal dye pairs were applied in the following order: (1) F4/80 at 1:500 dilution overnight at 4°C and Opal 570 at 1:150 dilution, (2) CD86 at 1:500 dilution for 30 minutes at room temperature and Opal 650 at 1:150 dilution, (3) CD206 at 1:400 dilution for 60 minutes at room temperature and Opal 520 at 1:150 dilution.

Whole slide scanned fluorescent images were evaluated to quantify cell infiltration and phenotype. Image deconvolution, annotations, threshold determination, and cell detection were performed using the open-source QuPath software package (v0.3.3). Annotations were manually created to include the whole scaffold and the scaffold area of visually dense cell infiltration. Trichrome images were used to define and quantify the scaffold boundary and area of full cell infiltration (Figure S7). These Trichrome annotations assisted in defining scaffold regions in the serially sectioned fluorescently stained slides. Fluorescent intensity thresholds to define positivity were determined by checking the pixel values and qualitatively cross-referencing whole slide scanned images (**SFigure 7**).

### 3.4. Quantitative Real-Time PCR

RNA from the harvested quadricep muscle was isolated using the RNeasy Mini Kit (Qiagen) according to the manufacturer’s protocol. 1 mL of Trizol was added to frozen tissue and homogenized using an OMNI GLH 850 Homogenizer in 10 second pulses. 0.2 mL of chloroform was added to the sample tube and shaken vigorously for 15 seconds, after which it incubated for 3 minutes at room temperature. Tubes were centrifuged at 12000 g for 15 minutes at 4 °C and the upper aqueous layer was removed and combined with 0.5 mL of 100% ethanol (Decon Labs). After incubating for 10 minutes at room temperature, up to 700 µL of the sample was loaded onto a RNeasy Mini spin column and centrifuged at 10000 g for 15 seconds. Flow-through was discarded and 80 µL of the DNase I incubation mix was added and incubated for 15 minutes at room temperature. 350 µL of RW1 buffer was added to the column, centrifuged at 10000 g for 15 seconds at room temperature, and flow-through was discarded. 500 µL of RPE buffer was added to the column, centrifuged at 10000 g for 30 seconds at room temperature, and flow-through was discarded. 500 µL of RPE buffer was added to the column, centrifuged at 10000 g for 2 minutes at room temperature, and flow-through was discarded. The RNA was eluted using 30-50 µL of RNase-free water after incubating for 10 minutes and centrifuging at 15000 g for 1 minute. RNA concentration and purity were confirmed using the Qubit RNA High Sensitivity Assay Kit and RNA Integrity and Quality Assay Kit (ThermoFisher Scientific). The Qubit RNA HS reagent and IQ reagent were diluted 1:200 in Qubit RNA HS buffer and IQ buffer respectively. Standards were prepared by mixing 10 µL of the corresponding standards in 190 µL of the working solution, vortexed, and incubated for 2 minutes at room temperature. 1-20 µL of the isolated RNA was added to 180-199 µL of the working solution for a final volume of 200 µL, vortexed, and incubated for 2 minutes at room temperature. RNA concentration and purity were calculated using a Qubit 4 Fluorometer (Invitrogen).

Up to 2.5 µg of the isolated RNA was reverse transcribed to cDNA using the SuperScript IV VILO Master Mix (Invitrogen) according to the manufacturer’s protocol. 4 µL of the SuperScript IV VILO Master Mix was combined with template RNA and nuclease-free water for a final volume of 20 µL. Using a T100 Thermal Cycler (Bio-Rad), the components incubated at 20 °C for 10 minutes to anneal primers, at 50 °C for 10 minutes to reverse transcribe RNA, and at 85 °C for 5 minutes to inactivate enzymes. The RNA and cDNA were isolated at -80 °C until use.

qRT-PCR was performed in triplicate to quantify the gene expression of Il4, Il10, Il17a, Il1b, Ccl9, and Il5 from cDNA using the FAM based TaqMan Gene Expression Assay (Applied Biosystems). 10 µL of TaqMan Gene Expression Master Mix, 1 µL of respective TaqMan Assay, and 9 µL of cDNA nuclease-free water were combined for a final 20 µL volume and loaded in a 96-well plate. The reaction plate was sealed with optical adhesive film and centrifuged to collect all the contents to the bottom. The plate was run in a Roche LightCycler 480 Instrument II and a Bio-Rad CFX Opus 96 Real-Time PCR System and programmed according to the manufacturer’s instruction. The fold change in gene expression was calculated using the 2^−ΔΔCt^ method.

### 3.5. Flow Cytometry

Tissue samples are prepared the same day as flow experiment. Freshly isolated quadriceps, spleens and lymph nodes were isolated and diced on ice into small pieces then digested using Liberase TL (0.25 mg/ml) and DNAse I (0.2 mg/ml) in 5mL RPMI for 25 min at 37°C with gentle shaking for spleen and lymph nodes and 45 minutes for quadriceps. The digestion mixture was passed through a 70µm cell strainer by grinding using a syringe plunger and cold PBS to rinse out cells. Single cells were centrifuged at 300g for 5 min at 4°C, and pelleted cells were washed with cold PBS. The cell pellet was resuspended in 5mL 1X RBC lysis buffer for 3 minutes on ice followed by two washes with cold PBS. A small portion of cells were stained with AO/PI for viability and cell counting.

We stained cells for viability using Zombie NIR for 25 minutes on ice in dark to exclude the dead cells followed by surface staining with antibody cocktail. 3X10^6^ cells from scaffold and spleen and 0.5X10^6^ cells from LN were stained for viability followed by antibody cocktail with the addition of Fc block and monocyte block in 100µL FACS buffer. The cells were incubated for 40 minutes on ice and washed with FACS buffer twice by centrifuging at 300g for 5 minutes at 4C. The surface-stained cells were fixed and permeabilized using Fix/Perm buffer for 25 minutes on ice followed by 2 washed with perm buffer. Next, intracellular staining was performed on fixed and permeabilized cells with the second antibody cocktail diluted in perm buffer for 30 minutes on ice followed by 3 washes with perm buffer and 1 wash with FACS buffer. Finally resuspended in 250µL FACS buffer and acquired on Cytek Aurora Spectral Flow Cytometer followed by analysis on SpectroFlo, OMIQ and FlowJo software.

### 3.6. Statistical Analyses

All statistical analyses were performed using GraphPad PRISM software (GraphPad Software, Boston, MA). All analyses were conducted using one-way ANOVA with posthoc Tukey test for pairwise comparisons. Significance is defined as P < 0.05, and is indicated as stated via indicator (*) or P-value.

## Acknowledgements

This Research was supported by the Center for Cancer Research, National Cancer Institute, National Institutes of Health Intramural Research Program project number ZIA BC 012021 and federal funds from the National Cancer Institute, National Institutes of Health, under contract HHSN261200800001E. The contributions of the NIH authors were made as part of their official duties as NIH federal employees, are in compliance with agency policy requirements, and are considered Works of the United States Government. However, the findings and conclusions presented in this paper are those of the authors and do not necessarily reflect the views of the NIH or the U.S. Department of Health and Human Services. The authors would like to thank Drs. Daniel McVicar, David Wink, Scott Durum, Stephen Anderson, and Leah Cook of the Cancer Innovation Lab at NCI, Frederick for their insightful discussions. We thank Megan Karwan and Jeff Carrell of the CCR-Frederick Flow Cytometry Core for cytometry support, the Laboratory Animal Sciences Program (LASP) for animal support, and the Frederick National Laboratory for Cancer Research NCI Frederick Molecular Histopathology Laboratory (MHL) for histology support.

## References

1 Kulwatno, J., Goldman, S. M. & Dearth, C. L. Volumetric Muscle Loss: A Bibliometric Analysis of a Decade of Progress. Tissue Eng Part B Rev 29, 299–309 (2023). 10.1089/ten.TEB.2022.0150

2 Frantz, C., Stewart, K. M. & Weaver, V. M. The extracellular matrix at a glance. J Cell Sci 123, 4195–4200 (2010). 10.1242/jcs.023820

3 Yang, D., Han, Z. & Oppenheim, J. J. Alarmins and immunity. Immunol Rev 280, 41–56 (2017). 10.1111/imr.12577

4 Yang, W. & Hu, P. Skeletal muscle regeneration is modulated by inflammation. J Orthop Translat 13, 25–32 (2018). 10.1016/j.jot.2018.01.002

5 Kurosaka, M., Hung, Y. L., Machida, S. & Kohda, K. IL-4 Signaling Promotes Myoblast Differentiation and Fusion by Enhancing the Expression of MyoD, Myogenin, and Myomerger. Cells 12 (2023). 10.3390/cells12091284

6 Wolf, M. T. et al. Polypropylene surgical mesh coated with extracellular matrix mitigates the host foreign body response. J Biomed Mater Res A 102, 234–246 (2014). 10.1002/jbm.a.34671

7 Daly, K. A. et al. The host response to endotoxin-contaminated dermal matrix. Tissue Eng Part A 18, 1293–1303 (2012). 10.1089/ten.TEA.2011.0597

8 Pal, S. et al. Extracellular Matrix Scaffold-Assisted Tumor Vaccines Induce Tumor Regression and Long-Term Immune Memory. Adv Mater 36, e2309843 (2024). 10.1002/adma.202309843

9 Wolf, M. T. et al. A biologic scaffold-associated type 2 immune microenvironment inhibits tumor formation and synergizes with checkpoint immunotherapy. Sci Transl Med 11 (2019). 10.1126/scitranslmed.aat7973

10 Lokwani, R. et al. Eosinophils Respond to Extracellular Matrix Treated Muscle Injuries but are Not Required for Macrophage Polarization. Adv Healthc Mater 14, e2400134 (2025). 10.1002/adhm.202400134

11 Sommerfeld, S. D. et al. Interleukin-36gamma-producing macrophages drive IL-17-mediated fibrosis. Sci Immunol 4 (2019). 10.1126/sciimmunol.aax4783

12 Lokwani, R. et al. Pro-regenerative biomaterials recruit immunoregulatory dendritic cells after traumatic injury. Nat Mater 23, 147–157 (2024). 10.1038/s41563-023-01689-9

13 Sadtler, K. et al. Developing a pro-regenerative biomaterial scaffold microenvironment requires T helper 2 cells. Science 352, 366–370 (2016). 10.1126/science.aad9272

14 Yang, Y. et al. Double-Negative T Cells Regulate Hepatic Stellate Cell Activation to Promote Liver Fibrosis Progression via NLRP3. Front Immunol 13, 857116 (2022). 10.3389/fimmu.2022.857116

15 Sicari, B. M. et al. A murine model of volumetric muscle loss and a regenerative medicine approach for tissue replacement. Tissue Eng Part A 18, 1941–1948 (2012). 10.1089/ten.TEA.2012.0475

