## Supplementary Information for "Inflammatory Agonists Modulate the Host Response to Type 2 ECM scaffold Immune Environment and Long-Term Remodeling After Severe Traumatic Injury"

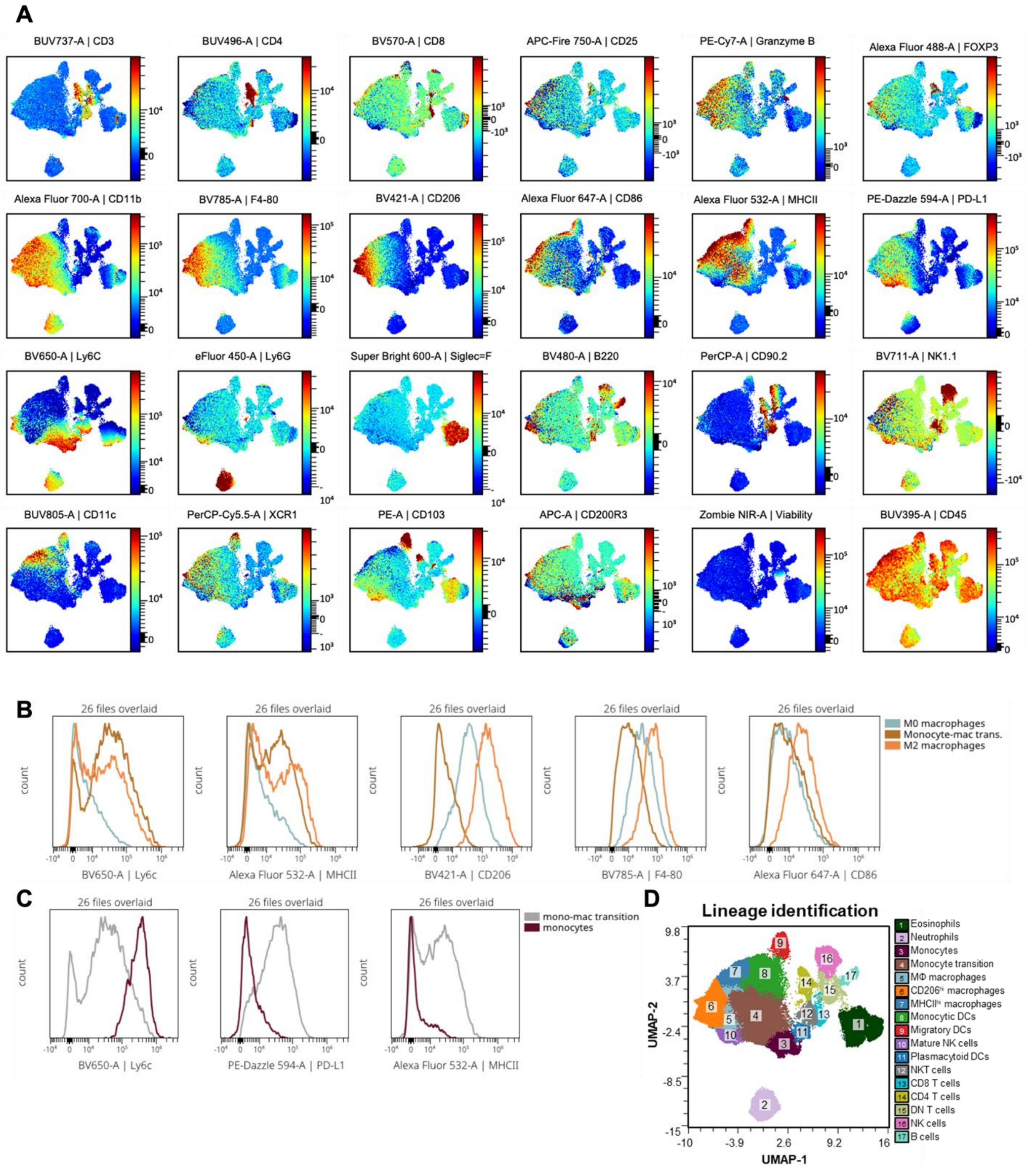


**SFigure 1: UMAP analyses. (A)** Week-1 UMAP with color coding the markers' expression level among all cells. Samples from all treatment groups were overlayed. **(B)** Histograms indicating the

expression profile differences among three macrophage subpopulations, M0, M2 and transitional macrophages. **(C)** Differentially expressed markers between monocytes and transitional macrophages. **(D)** Meta clustered UMAP.

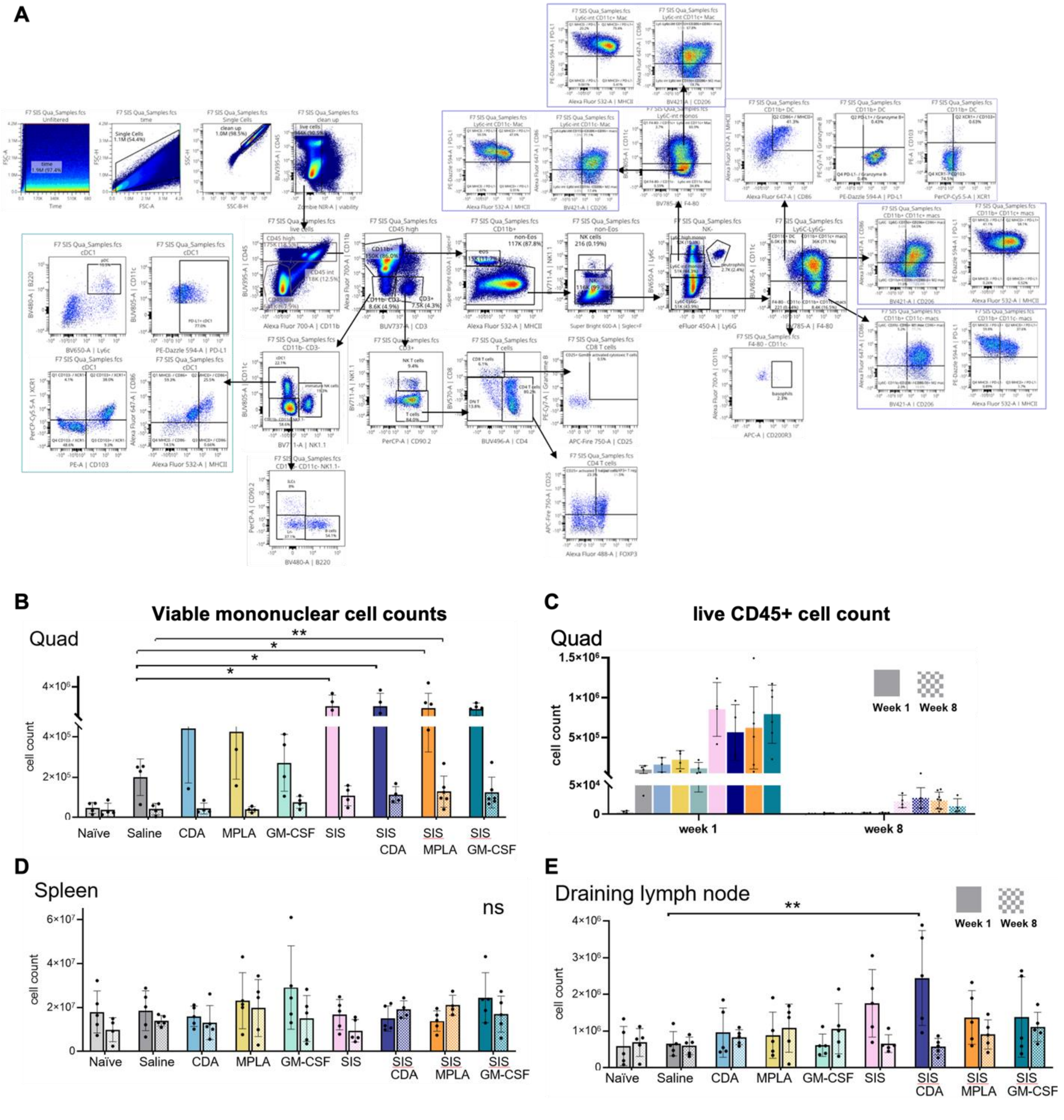

**SFigure 2: (A)** Flow cytometry gating strategy. Viable mononuclear cell counts from AOPI staining after tissue harvesting from **(B)** quadricep, **(D)** spleen and **(E)** draining lymph node at week 1 and 8. **(C)** Live CD45+ cell count from flow cytometer at week 1 and 8.

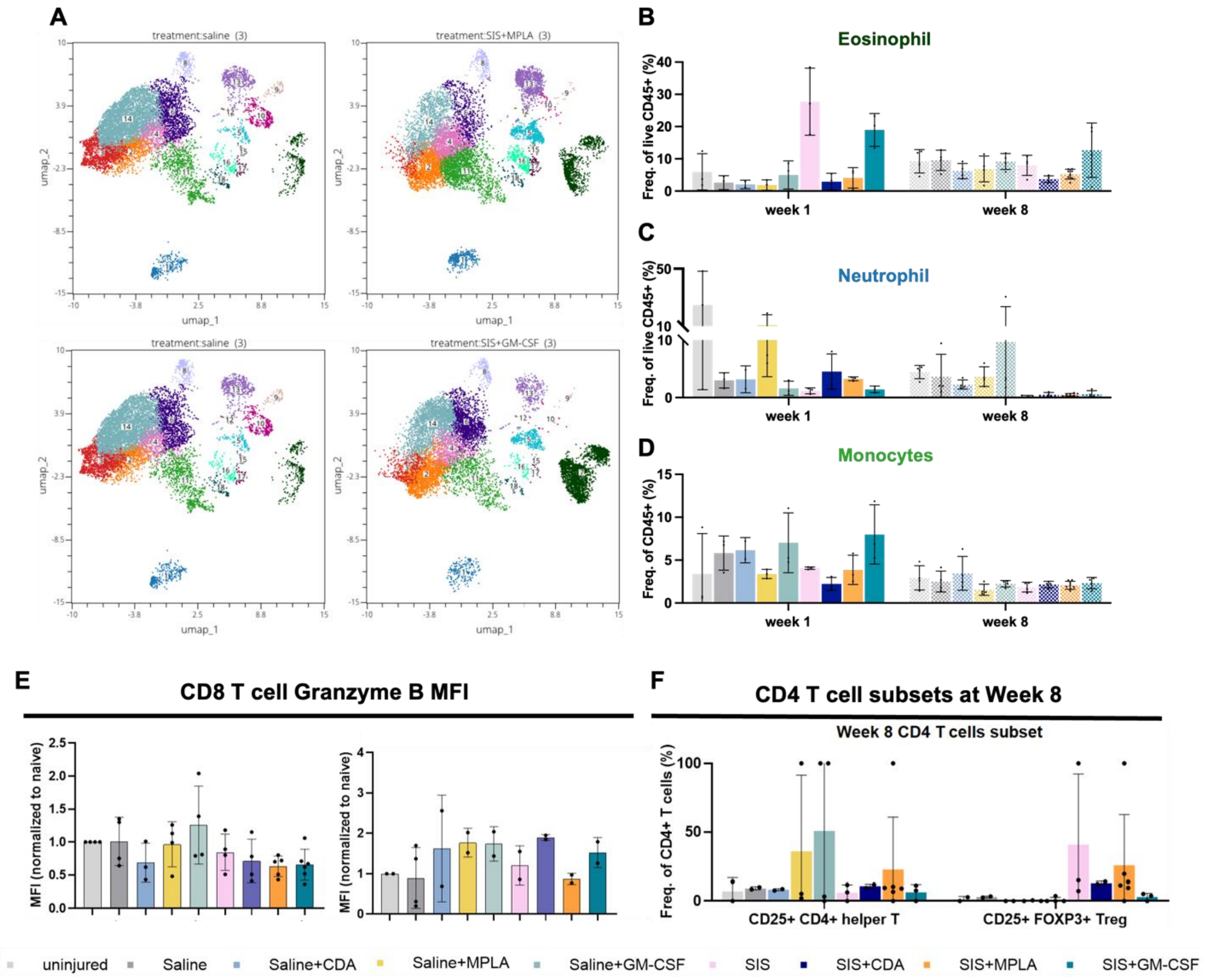


**SFigure 3: (A)** Clustered UMAP comparison among treatment groups. Percentage comparison of **(B)** eosinophils, **(C)** neutrophils and **(D)** monocytes among tr treatment groups at week 1 and 8. **(E)** Granzyme B mean fluorescent intensity comparison of CD8 T cells among treatment groups at week 1 and 8. **(F)** CD4 T cell activation state comparison at week 8.

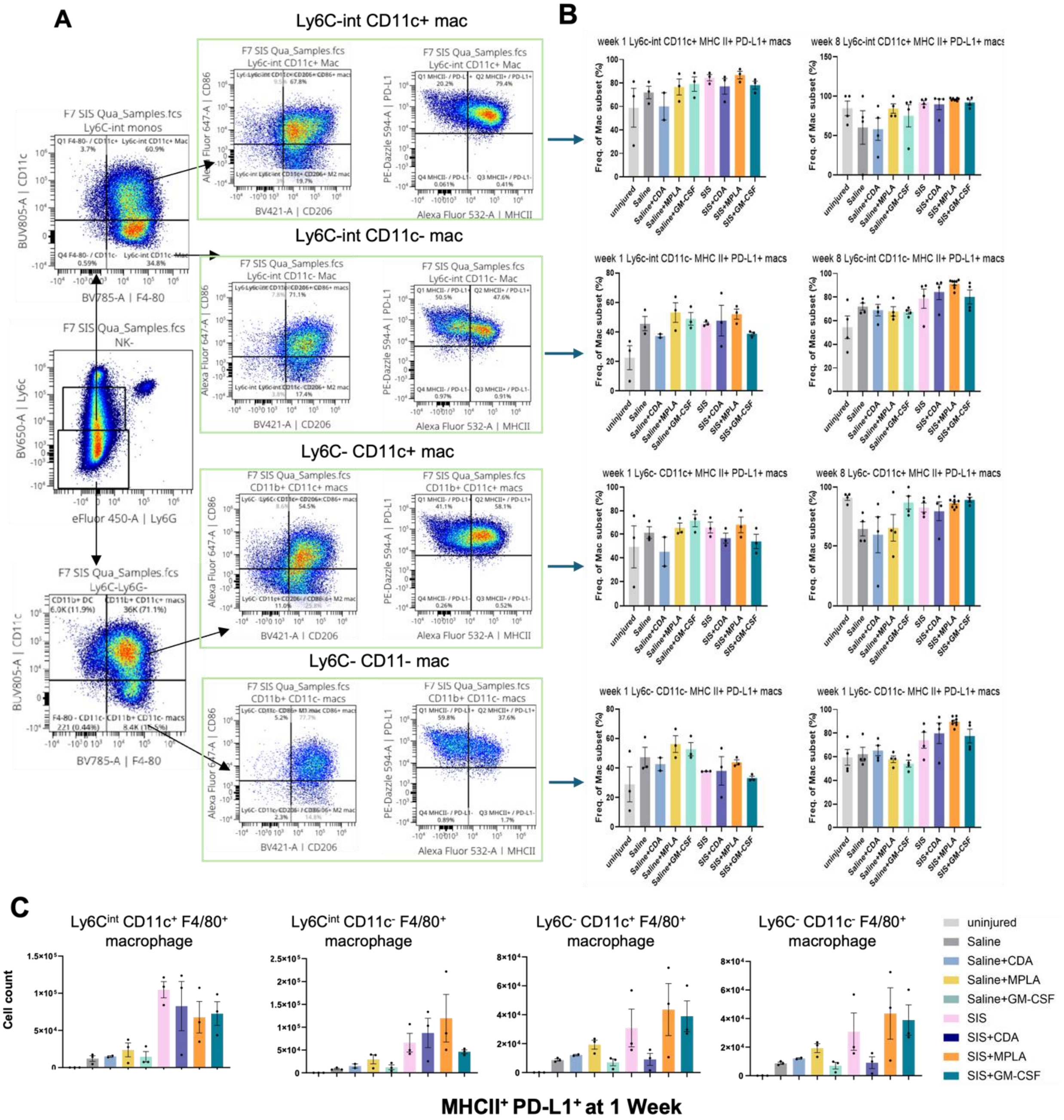


**SFigure 4: (A)** Macrophage subsets gating. **(B)** Macrophage subsets comparison in MHCII and PD-L1 expression levels at week 1 and 8. Comparison was made as percentage of macrophage subset. **(C)** Macrophage subsets cell count comparison at week 1.

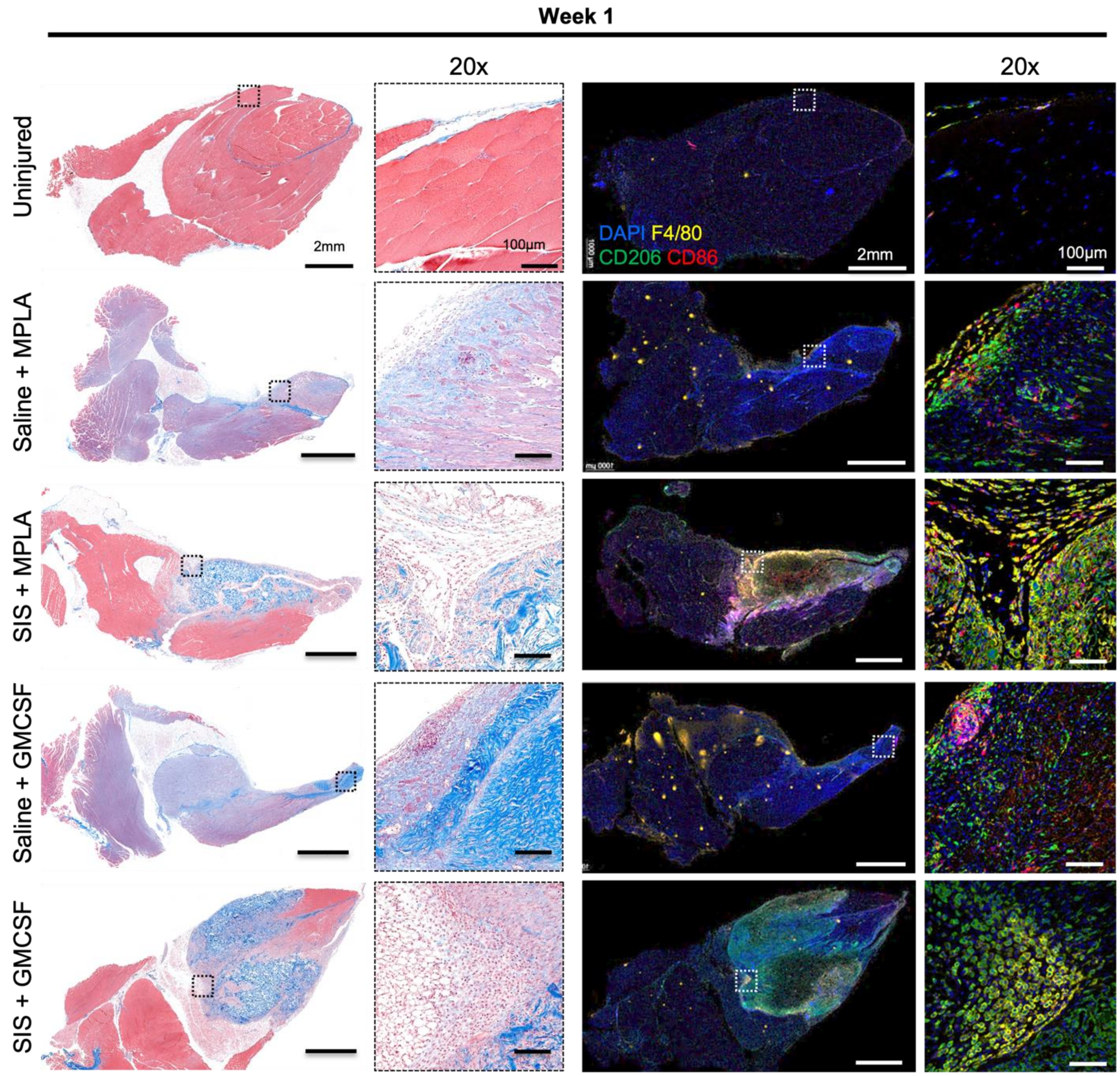


**SFigure 5: Histologic and spatiotemporal immune profiling of VML injury post week 1 of ECM scaffold implantation.** Masson's Trichrome-stained images (Left panel) and multiplex immunofluorescent images (Right panel) showing the morphology of VML injury microenvironment with immune agonist co-deliver (MPLA, and GM-CSF) with SIS-ECM scaffold or saline post 1 week of injury and implantation in C57Bl/6 mice quadriceps. Boxes highlight immune infiltrates at the at 20X objective.

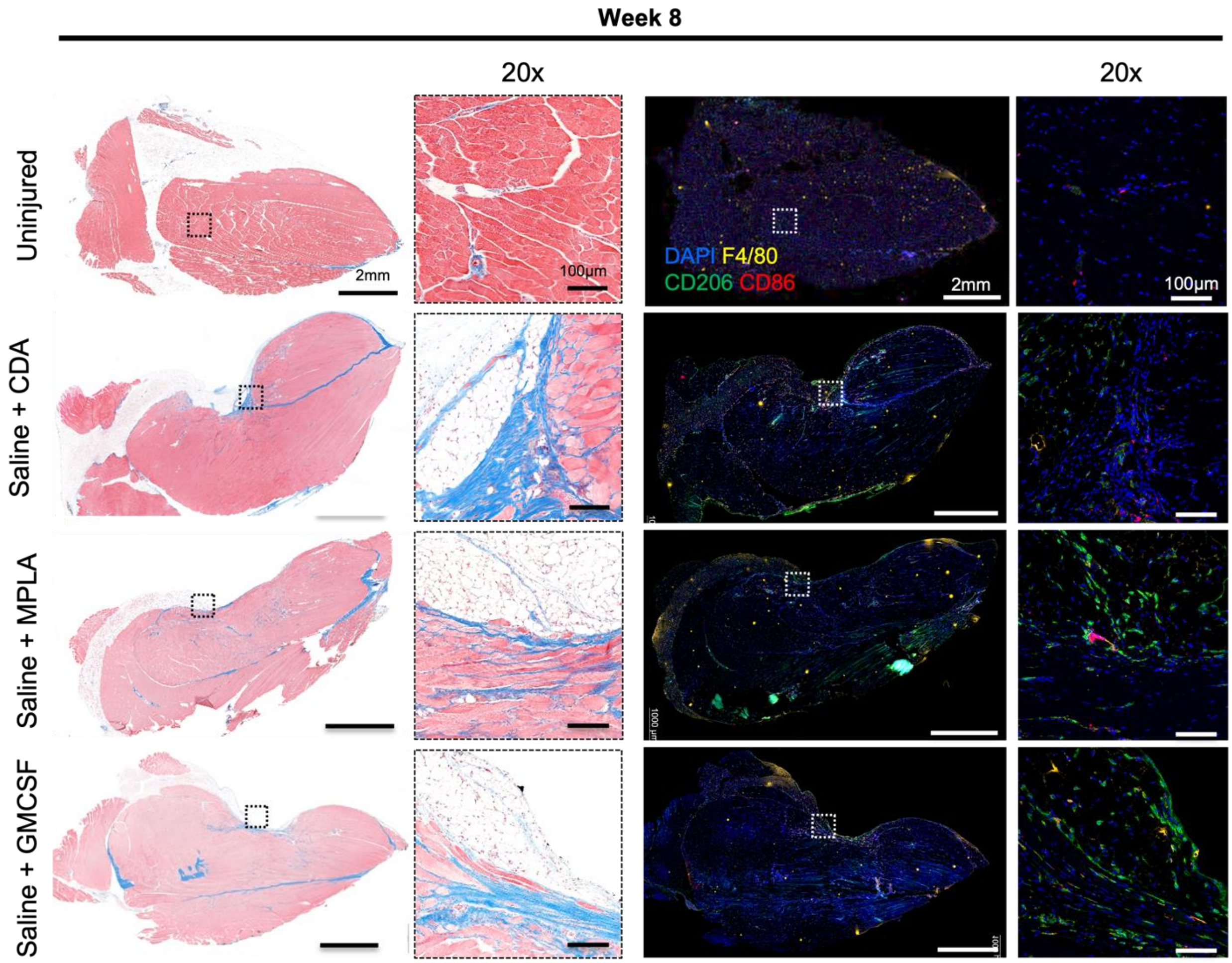


**SFigure 6: Histologic and spatiotemporal immune profiling of VML injury post week 8 of ECM scaffold implantation.** Masson's Trichrome-stained images (Left panel) and multiplex immunofluorescent images (Right panel) showing the morphology of VML injury microenvironment with immune agonist co-deliver (CDA, MPLA, or GM-CSF) with saline post 8 weeks of injury and implantation in C57Bl/6 mice quadriceps. Boxes highlight immune infiltrates at the at 20X objective.

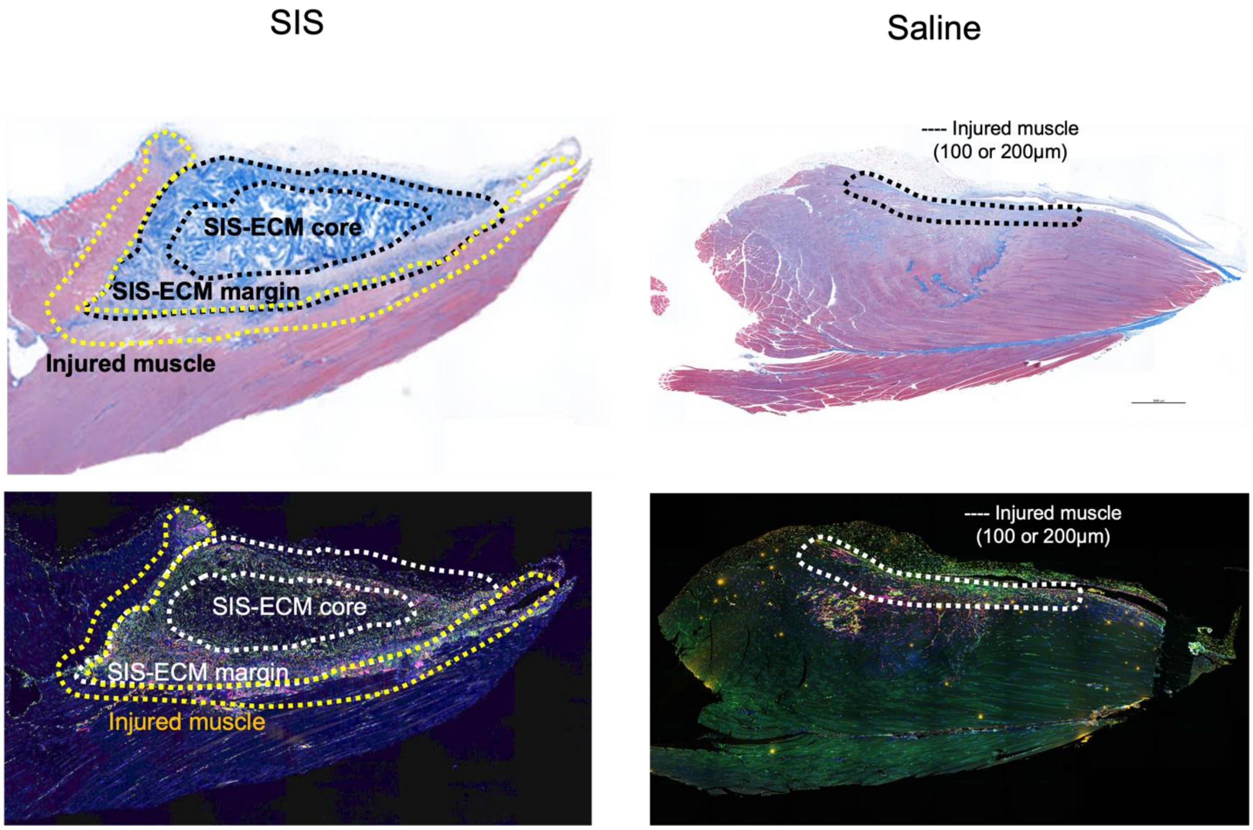


**SFigure 7: Schematics showing the image annotation for cell quantification.** Masson's Trichrome-stained images and multiplex immunofluorescent images showing the annotation used for quantification of immune cells in different region of the quadriceps muscle post injury and implantation with SIS-ECM scaffold or saline. The saline group has only injured muscle region (200µm from injury site towards the muscle) whereas the SIS-ECM scaffold group has three regions as marked in the image above showing the injured muscle region followed by SIS-ECM margin (200µm from injury site towards ECM) and there is a SIS-ECM core at the center of the scaffold.

**Table 1: Antibodies used with dilution, clone no., Cat# and Company.**

| Target | Conjugate | Clone | species | Dilution | Company, Cat # |
|---|---|---|---|---|---|
| CD206/MRC1 | - | E6T5J | rabbit | 1:400 | Cell Signaling Technology, Cat# 24595T |
| F4/80 | - | BM8 | rat | 1:500 | BioLegend, Cat# 123101 |
| CD86 | - | E5W6H | rabbit | 1:500 | Cell Signaling Technology, Cat# 20018 |
| Anti-rabbit IgG | HRP polymer | - | - | Neat | Biocare Medical, REF# RMR622H |
| Anti-rat IgG | HRP polymer | - | - | Neat | Biocare Medical, REF# BRR4016H |

**Table 2: Antibodies used with dilution, clone no., Cat# and Company.**

| Fluorophore | Wavelength (nm) | Dilution | Manufacturer, Catalog # |
|---|---|---|---|
| Opal 570 | 550/570 | 1:150 | Akoya Biosciences, #FP1488001KT |
| Opal 650 | 627/650 | 1:500 | Akoya Biosciences, #FP1496001KT |
| Opal 520 | 494/525 | 1:150 | Akoya Biosciences, #FP1487001KT |
